# Drosophila Mechanical Nociceptors Preferentially Sense Localized Poking

**DOI:** 10.1101/2022.01.04.474910

**Authors:** Zhen Liu, Qi-Xuan Wang, Meng-Hua Wu, Shao-Zhen Lin, Xi-Qiao Feng, Bo Li, Xin Liang

## Abstract

Mechanical nociception is an evolutionarily conserved sensory process required for the survival of living organisms. Previous studies have revealed much about the neural circuits and key sensory molecules in mechanical nociception, but the cellular mechanisms adopted by nociceptors in force detection remain elusive. To address this issue, we study the mechanosensation of a fly larval nociceptor (class IV da neurons, c4da) using a customized mechanical device. We find that c4da are sensitive to mN-scale forces and make uniform responses to the forces applied at different dendritic regions. Moreover, c4da showed a greater sensitivity to more localized forces, consistent with them being able to sense the poking of sharp objects, such as wasp ovipositor. Further analysis reveals that high morphological complexity, mechanosensitivity to lateral tension and active signal propagation in the dendrites altogether facilitate the mechanosensitivity and sensory features of c4da. In particular, we discover that Piezo and Ppk1/Ppk26, two key mechanosensory molecules, make differential but additive contributions to the mechanosensation of c4da. In all, our results provide updates into understanding how c4da process mechanical signals at the cellular level and reveal the contributions of key molecules.

## Introduction

Mechanosensation is a physiological process that transduces mechanical stimuli into neural signals (Chalfie, 2009; Marshall and Lumpkin, 2012). It underlies the perception of gentle touch, sound, acceleration and noxious force. To cope with manifold environmental forces, mechanoreceptor cells are diversified (Lumpkin et al., 2010; Zimmerman et al., 2014). Among different types of mechanoreceptor cells, those activated by noxious forces, i.e. mechanical nociceptors, are of particular importance because they are essential for the survival of living organisms (Lumpkin et al., 2010; Tracey, 2017).

Much effort has been made to understand the mechanical nociception in several model organisms. In mammals, free nerve endings of nociceptive neurons penetrate the keratinocyte layer of skin and serve as the primary nociceptors. The axons of these neurons output to neural circuits in the spinal cord, which then transmit pain signals to local interneurons or up to the brain to initiate neural reflexes (Tracey, 2017). At the molecular level, transient receptor potential (TRP) channels, Piezo and other channels are found to be involved in mechanical nociception (Kwan et al., 2006; Murthy et al., 2018; Tracey, 2017). In lower animals (such as worms and flies), mechanical nociception has also been extensively studied (Chatzigeorgiou et al., 2010; Tracey, 2017). A widely used model is fly larval mechanical nociception mediated by class IV dendritic arborization neurons (c4da). C4da are polymodal nociceptors that can be activated by light, thermal and mechanical (e.g. harsh touch) stimuli (Hwang et al., 2007; Kim et al., 2012; Terada et al., 2016; Xiang et al., 2010). The axon of c4da synapses to several targets in the ventral nerve cord, which then output signals to trigger the rolling behavior, a stereotyped locomotion in escaping from nociceptive stimuli (Burgos et al., 2018; Grueber et al., 2007; Yoshino et al., 2017). Furthermore, two parallel pathways in c4da, one mediated by Ppk1/Ppk26 and the other by Piezo, are required for the behavioral responses in mechanical but not thermal nociception (Gorczyca et al., 2014; Guo et al., 2014; Kim et al., 2012; Mauthner et al., 2014).

Despite the known information about the neural circuits and sensory molecules in the c4da-mediated mechanical nociception, the cellular mechanism of how c4da detect forces remains unclear. A recent study shows that the nociceptive responses of c4da to thermal stimuli depend on the dendritic calcium influx through two TRPA channels and the L-type voltage-gated calcium channel, suggesting the presence of neuronal processing of heat-induced responses (Terada et al., 2016). In a more general sense, it is also intriguing how mechanoreceptor cells are optimized for their specific stimuli. An interesting study reports unique neuronal mechanisms in the mechanosensory neurons in tactile-foraging birds (Schneider et al., 2014), also demonstrating the presence of cellular mechanisms that facilitate the response to specific stimuli. So far, how c4da process mechanical inputs in nociception at the neuronal level remains unclear.

To address this issue, we build a “mechanical-optical” recording system that is able to measure the in vivo sensory response of c4da to the stimulating forces with controlled strength, contact area and force application point. We find that c4da are sensitive to mN-scale force, and moreover, c4da use their entire dendritic territory as the force-receptive field. In particular, c4da show greater responses to the stimuli delivered using small probes, suggesting their preferential sensitivity to localized forces. Further analysis reveals the cellular mechanisms that facilitate the sensory features of c4da and examines the contributions of key molecules. These findings update the current model for the c4da-mediated mechanical nociception and provide mechanistic insights into how mechano-nociceptor cells process force signals at the cellular level.

## Results

### Mechanical recording and the “sphere-surface” contact model

We set out by building a mechanical device that was able to exert and measure compressive forces on fly larval fillets (Fig. 1A). This device contained three parts: a piezo stack actuator, a metal beam coupled with a strain gauge and a force probe made of capillary glass tube (Fig. 1A and Fig. s1). The whole device was mounted on the working stage of an inverted spinning-disk confocal microscope (Fig. 1A and Fig. s1). To record the force-evoked response of sensory neurons, freshly prepared larval fillet was spread and mounted on a polydimethylsiloxane (PDMS) pad (thickness: 1 mm) with the exterior surface of the cuticle accessible to the force probe and the interior surface visible to the confocal microscope (Fig. 1A). When the force probe, driven by the piezo actuator, delivered compressive forces onto the larval fillet, the strain gauge converted the deflection of the metal beam into an electrical signal. This electrical signal could be translated into a force signal using the calibration curve (Fig. 1 B-C, see Methods). By making spherical probes in different diameters (Fig. 1D), we can study how mechanoreceptor cells respond to the probes of different sizes. In the meantime, we also monitored the position of force probes, dendritic morphology (membrane marker) and neuronal response (GCaMP6s) (Fig. 1E).

**Figure 1.**
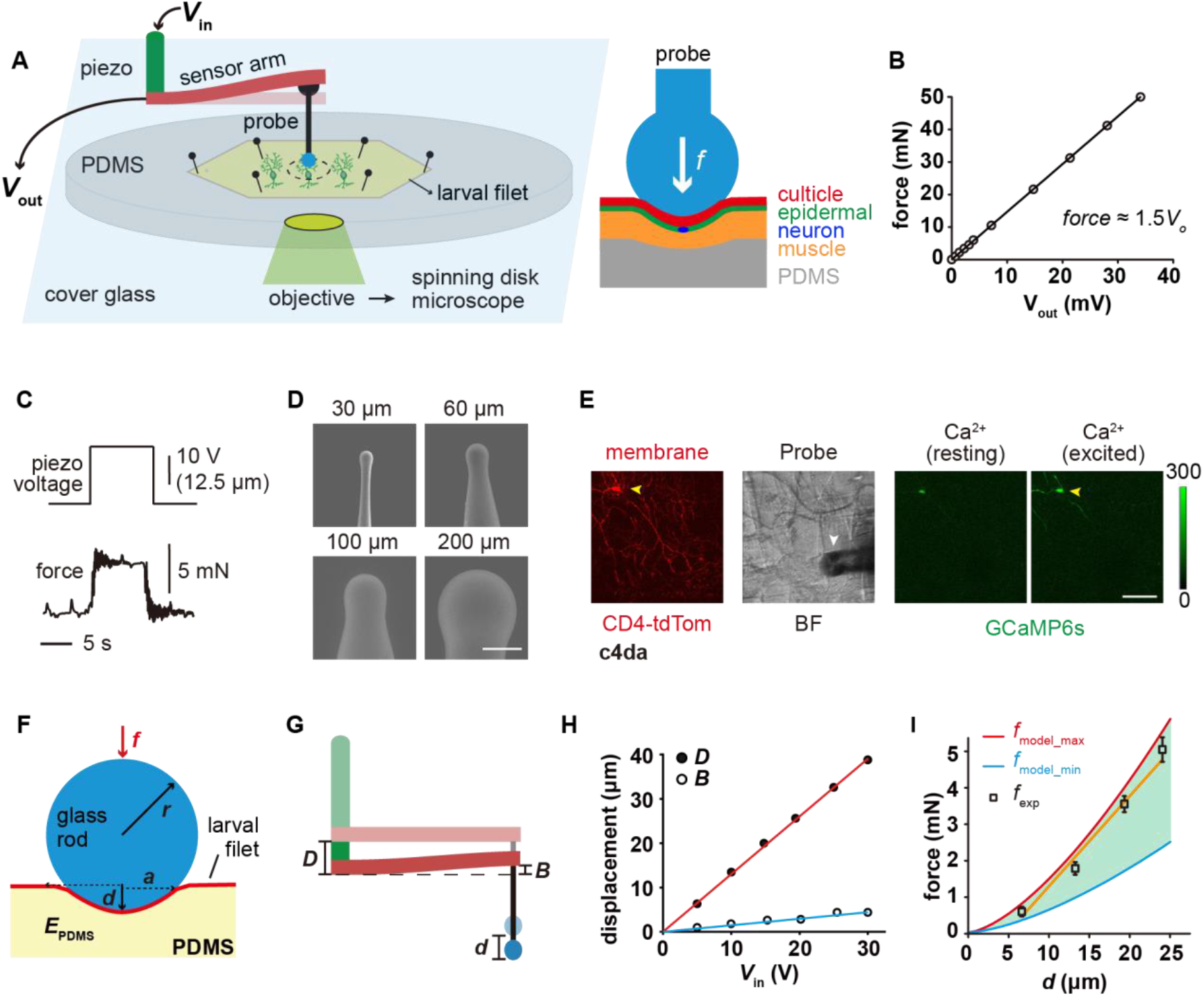
The “mechanical-optical” recording system. **(A)** The cartoon schematics for the “mechanical-optical” recording system (left) and the contact model between a spherical probe and larval fillet (right). *V*_in_ was the driving voltage of the piezo actuator. *V*_out_ was the readout voltage of the strain gauge. **(B)** The force calibration curve of the strain gauge. The data points were mean values from three measurements. **(C)** Representative traces for the input (driving voltage of the piezo actuator, upper panel) and the mechanical output (force, lower panel) of the recording system. **(D)** The scanning electron microscopy images of glass force probes of different sizes. Scale bar, 100 μm. **(E)** Left panel: a representative image of c4da (membrane, red channel), Middle panel: a bright field image of larval fillet. Right panel: representative images showing GCaMP6s signals in c4da at resting (left) and exciting (right) conditions. Yellow arrowhead: soma. White arrowhead: force probe. Scale bar, 100 μm. Genotype: *uas-cd4-tdTom*; *ppk-gal4/20×uas-ivs-gcamp6s*. **(F)** The mechanical schematics of the “sphere-surface” model. Note that the contact interface had a spherical crown shape. The definitions of all model parameters were described in Methods. **(G)** The cartoons schematics showing the relationship among indentation depth of the force probe (*d*), deflection of the beam (*B*) and stepping distance of the piezo (*D*). **(H)** The plots of *D* (red) and *B* (blue) versus the driving voltage of the piezo (*V*_in_). **(I)** The comparison between the calculated (red and blue) and experimentally measured (black) contact forces. In our calculations, the maximal (4 MPa, red) and minimal (1.6 MPa, blue) values of the elastic modules of PDMS were from literatures (Sharfeddin et al., 2015; Vlassov et al., 2018). Data were presented as mean±std (*n*=9 assays).

The contact mechanics between the force probe and fillet can be approximated using a classic “sphere-surface” contact model (Johnson, 1985), which would allow us to distinguish the effects of different parameters, such as pressure and contact area. Note that to guarantee the accuracy of the contact model, the indentation depth (*d*) needs to be smaller than the radius of the spherical probe (*r*) (Fig. 1F). This prerequisite is acceptable for our measurements because pilot experiments showed that when the step distance (*D*) of the piezo actuator was greater than *r*, the force probe often penetrated the cuticle and caused tissue damage. This would interfere with our focus on mechanical nociception by introducing potential effects of chemo-nociception. To confirm if the modeling approximation is valid, we calculated the contact force (*f*_model_) from *d* (**Eq. 1** and **2**, see Methods), which could be obtained from the measured values of *D* and beam deflection (*B*) (**Eq. 3**, see Methods). *f*_model_ was then compared to the experimentally measured forces (*f*_exp_) (Fig. 1 G-I). As shown in Fig. 1I, the relationship between *f*_exp_ and *d* fell in the range of model calculations, demonstrating that the model approximation is valid. Note that different values for the elastic modulus of PDMS (*E*_PDMS_) (Sharfeddin et al., 2015; Vlassov et al., 2018) were used to calculate the upper and lower bounds of contact forces (Fig. 1I).

### C4da are sensitive to mN-scale forces and more sensitive to small probes

Using the customized mechanical device, we applied poking forces onto the c4da at the positions about 100 μm away from the soma (i.e. the “proximal” region, denoted as ***p*** in the figures) (Fig. 2A). By simultaneously monitoring probe location and neuronal morphology, we ensured that the probe made direct compression onto the dendrites. Force titration experiments using a 60 μm probe showed the half-activation force (*f*_50_) of ∼3 mN and the full-activation force (*f*_90_) of ∼4 mN, while those using a 30 μm probe showed the *f*_50_ of ∼0.7 mN and the *f*_90_ of ∼1.5 mN (Fig. 2 B-C), suggesting that c4da have different sensitivities to different probe sizes. To explore the entire dendritic field, we then compressed the dendrites of c4da at the positions about 200 μm away from the soma (i.e. the “distal” region, denoted as ***d*** in the figures) (Fig. 2D). In the condition of using a 60 μm probe and the saturating force (4 mN), the responses of c4da were not significantly different from those to the proximal stimuli (Fig. 2E). In the case of using a 30 μm probe and also a saturating force (1.5 mN), the responses of c4da were slightly weaker than those to the proximal stimuli, but most of the neurons did make clear responses (Fig. 2E). Therefore, our results showed that c4da use their entire dendritic field as the force-receptive field.

**Figure 2.**
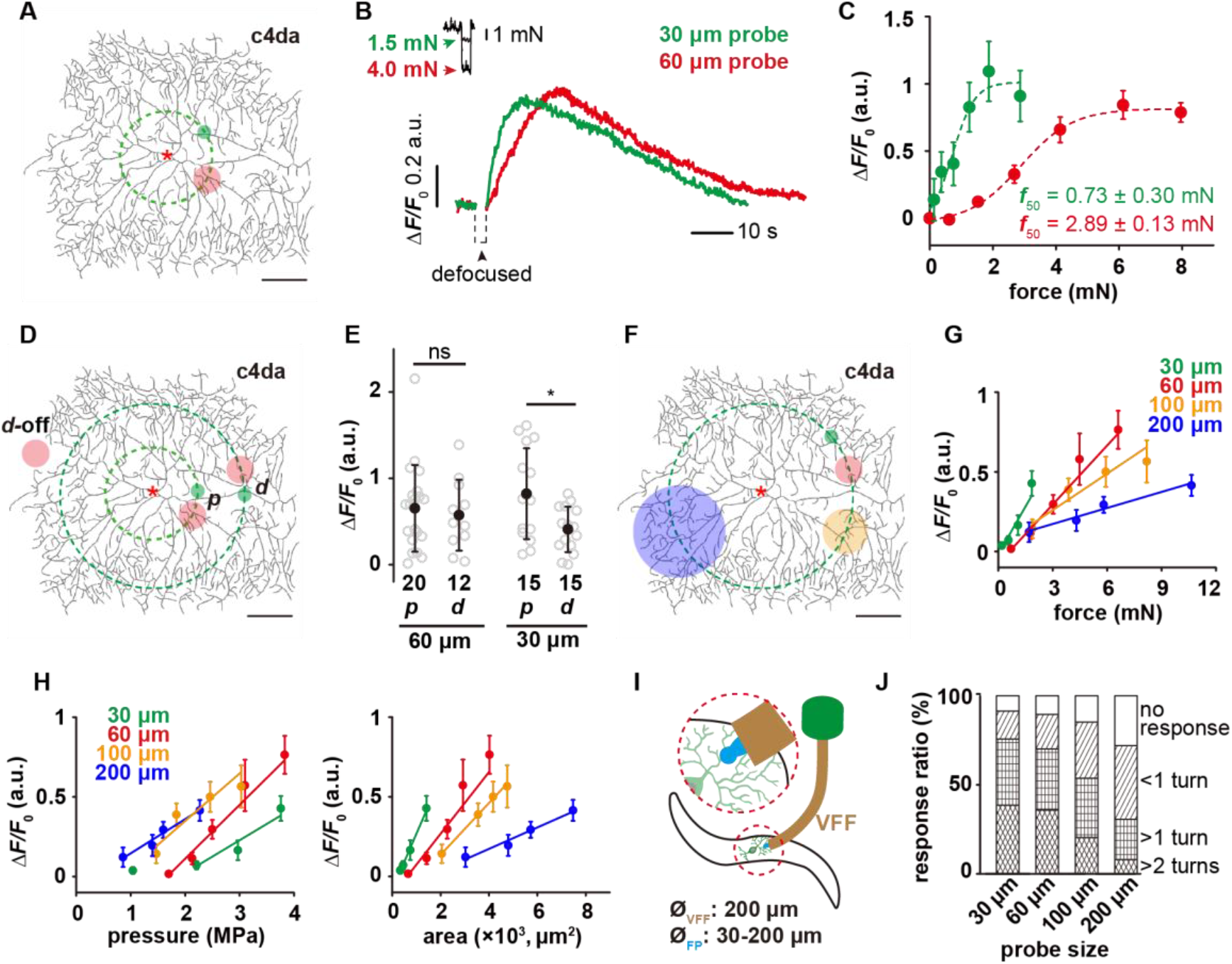
The mechanosensory features of c4da. **(A)** A representative image of c4da. The forces were applied at about 100 μm from the soma, i.e. along the green dashed circle. The representative force application points were marked using the filled circles (Green: 30 μm probe. Red: 60 μm probe). Genotype: *uas-cd4-tdtom*; *ppk-gal4*. **(B)** Representative responses of c4da (Δ*F*/*F*_0_, i.e. the change in calcium signal) to mN-scale forces delivered using the 30 μm (green) and 60 μm (red) probes. The black arrowhead indicated the defocused period of the soma caused by the stimulating force (2 s). Genotype: *ppk-gal4/+*; *ppk-cd4-tdtom/20×uas-ivs-gcamp6s*. **(C)** The force-response (Δ*F*/*F*_0_) plots of c4da (*n*=12 cells) at two different probe sizes. The dashed lines were Boltzmann fitting. **(D)** The schematic showing the force application points (filled circles, green: 30 μm probe, red: 60 μm probe) of different stimuli. The dashed concentric circles were 100 and 200 μm in radius, respectively. ***p***: proximal dendrite. ***d***: distal dendrite. ***d*-off**: the “dendrite-off” region. **(E)** The responses of c4da to the proximal and distal stimuli using the 30 (1.5 mN) and 60 μm (4 mN) probes. **(F)** The schematic showing the stimuli (filled circles, green: 30 μm probe, red: 60 μm probe, orange: 100 μm probe, blue: 200 μm probe) delivered using the probes of different sizes. The dashed circle was 200 μm in radius. **(G)** The responses of c4da (*n*=10 cells) to the forces applied on distal dendrites using the probes of different sizes. **(H)** The plots of the responses of c4da to distal stimuli versus central pressure (*P*_0_) (left panel) and contact areas (*A*_c_) (right panel), respectively. Also see Fig. s3 for the plots of the responses versus the pressures at other positions (*P*_x μm_). **(I)** The cartoon schematics for the modified behavior assay for mechanical nociception. VFF: Von Frey fiber, FP: force probe. **(G)** The behavioral responses of wild-type larvae to the mechanical poking stimuli using the probes of different sizes. 30 μm probe (*n* = 87 larvae), 60 μm probe (*n* = 102 larvae), 100 μm probe (*n* = 92 larvae), 200 μm probe (*n* = 72 larvae). In panels **A**, **D** and **F**, scale bar: 100 μm. In panels **C**, **G** and **H**, data were presented as mean±sem. In panel **E**, data were presented as mean±std and the numbers of cells were indicated below the scattered data points. Asterisk: the soma. *: p<0.05. ns: no significance.

To further justify the validity of this assay, we performed the same assay on c1da (class I da neuron) and c3da (class III da neuron). C1da showed no responses to compressive forces (Fig. s2), in consistence with their function being a proprioceptor rather than a tactile receptor (Guo et al., 2016). C3da showed a much smaller threshold of activation, suggesting a higher sensitivity in detecting tactile forces (using both 30 and 60 μm probes). This is consistent with their known function as the larval gentle touch receptor (Fig. s2) (Yan et al., 2013).

The observation of different responses of c4da to 30 and 60 μm probes suggests that c4da may be more sensitive to smaller probes. To test this idea, we stimulated c4da using probes of different diameters (30, 60, 100 and 200 μm) at the distal regions (Fig. 2F). At the same force, c4da made stronger responses (Δ*F*/*F*_0_) to smaller probes (Fig. 2G). Furthermore, the slope of the force-Δ*F*/*F*_0_ curve increased as the probe became smaller (Fig. 2G), reflecting an enhanced mechanosensitivity to the change in stimulating force. To further understand this result, we calculated central pressure (*P*_0_) and contact area (*A*_c_) based on the “sphere-surface” model (**Eq. 4-7**, see Methods), and then plotted the responses against these two parameters separately (Fig. 2H). In the *P*_0_-Δ*F*/*F*_0_ plots (left panel in Fig. 2H), we found that increasing contact area decreased the threshold of cell activation but had only minor impact on the slope of the curves, suggesting that c4da have a robust sensitivity to the change in pressure. Similar observations were obtained when the pressures at other positions (*P*_x μm_) were plotted against the responses (Δ*F*/*F*_0_) (Fig. s3). In the *A*_c_-Δ*F*/*F*_0_ plots (right panel in Fig. 2H), when the contact area was the same, higher pressure led to stronger responses (Fig. 2H), also in consistence with c4da being a pressure sensor. The comparison of the Δ*P* (the change of pressure) and Δ*A*_c_ (the change of contact area) of different probes showed that for a given change of forces, the smaller probes caused a smaller Δ*A*_c_ but a greater Δ*P* in comparison to the larger probes (Fig. 2H), which explains the higher sensitivity of c4da to sharper probes.

To further understand how the preferential sensitivity of c4da to localized force contributes to larval responses in nociception, we performed the behavior assay of mechanical nociception using a modified Von Frey fiber. To test the effect of different probe sizes, we attached the glass probes (shorter than 1 mm) of different diameters (from 30 to 200 μm) to one end of a Von Frey fiber (Fig. 2I). These modified fibers were used to poke fly larvae with a stimulating force of 10-20 mN. The defensive rolling behaviors were recorded and graded to reflect the sensory response of c4da. We found that in general, the larvae showed stronger behavioral responses to smaller probes (Fig. 2G), in consistence with our finding that c4da make stronger cellular responses to smaller probes.

In summary, our results demonstrate that c4da are sensitive to mN-scale compressive forces and use their entire dendritic arbor as the force receptive field. In particular, c4da are more sensitive to forces delivered using the small probes, consistent with c4da acting as nociceptors to sense mechanical poking or sting from sharp objects, e.g. the ovipositor of wasp (Cerkvenik et al., 2017; Hwang et al., 2007).

### Morphological complexity contributes to the sensory features of c4da

Having characterized the mechanosensory responses of c4da, we wondered what could be the cellular mechanisms underlying the sensory features of c4da, in particular the preferential sensitivity to small probes? The key thing is to ensure that an adequate number of mechanosensory molecules can be activated, even in the dendrites in distal regions or under a localized stimulation. Intuitively, a dense and uniform distribution of dendrites might facilitate these sensory features. We tested this hypothesis by studying the sensory responses of a knock-down mutant of *cut* (*ct*^i^), a key transcription factor that is known to determine the high complexity of c4da (Jinushi-Nakao et al., 2007; Sulkowski et al., 2011).

C4da-*ct*^i^ neurons showed a much sparser dendritic morphology than c4da-*wt* (Fig. 3A). Modified Sholl analysis showed that the dendritic density of c4da-*wt* was uniform across the entire dendritic field, while that of c4da-*ct*^i^ was similar to c4da-*wt* in the proximal region but rapidly decreased towards the distal region (Fig. 3B). In comparison to c4da-*wt* or c4da-scrambled RNAi, c4da-*ct*^i^ showed similar responses to the stimuli on proximal dendrites, but significantly weaker responses to those on distal dendrites (Fig. 3C). These results show that with a reduced morphological complexity, c4da-*ct*^i^ cannot use its entire dendritic arbor as an effective force receptive field. We also studied the responses of c4da-*ct*^i^ to the probes of different sizes. Although c4da-*ct*^i^ still showed force-dependent sensory responses, the overall response amplitudes were significantly reduced. In particular, the cellular responses to the 30 μm probe were reduced to the greatest extent (Fig. 3D). In consistence with the cellular observations, the defensive rolling behaviors of c4da-*ct*^i^ in response to the mechanical poking of small probes (30 and 60 μm) were largely weakened in comparison to those of c4da-*wt*, while those to large probes (e.g. 100 μm) showed nearly no change (Fig. 3E). Based on these observations, we conclude that the sensory preference of c4da to small probes was lost in c4da-*ct*^i^.

**Figure 3.**
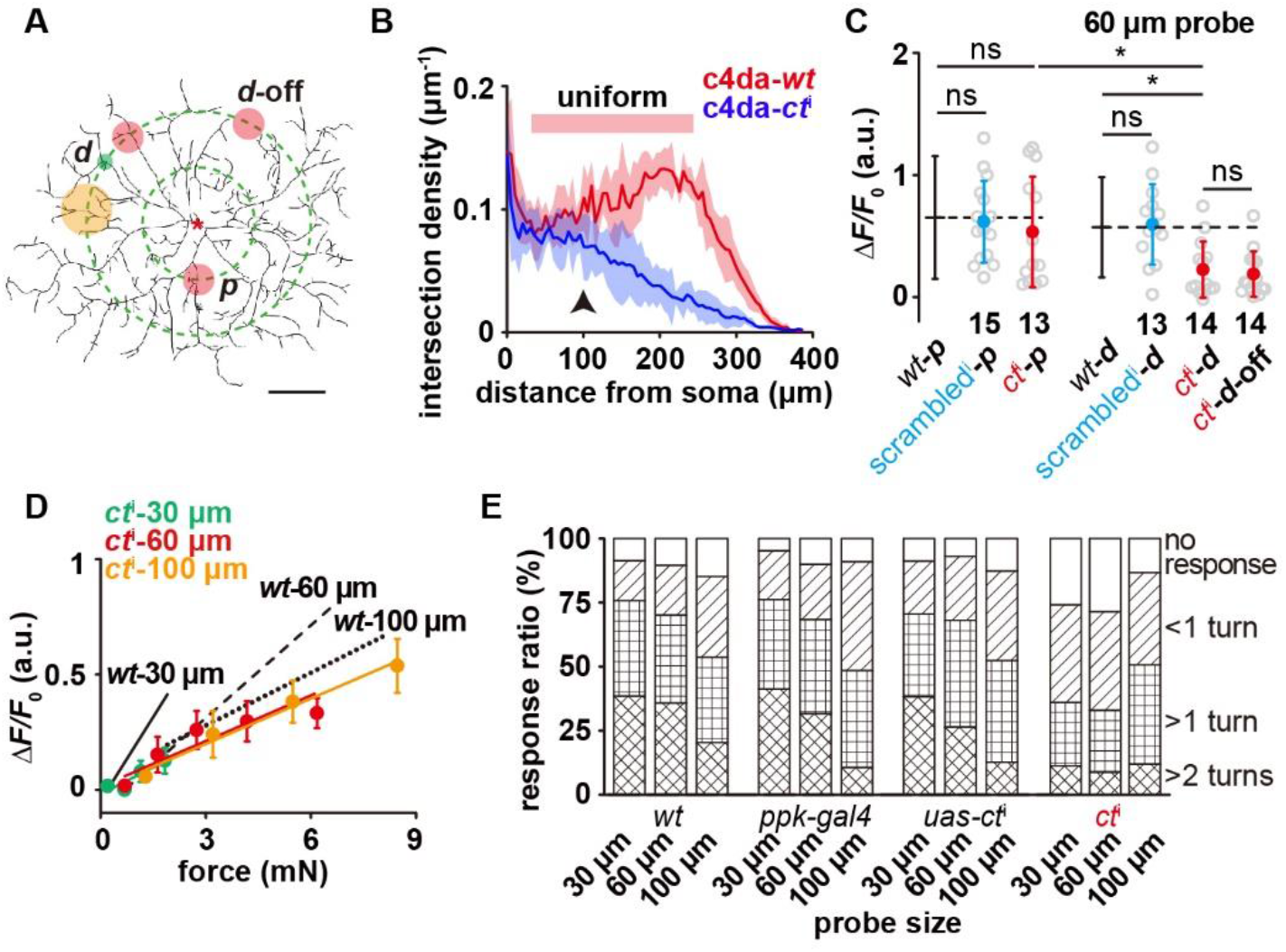
The contribution of dendritic morphology to the sensory features of c4da. **(A)**The schematic showing the force application points (filled circles, green: 30 μm probe, red: 60 μm probe, orange: 100 μm probe) on c4da-*ct*^i^. The dashed concentric circles were 100 and 200 μm in radius, respectively. ***p***: proximal dendrite. ***d***: distal dendrite. ***d*-off**: the “dendrite-off” region. Asterisk: the soma. Scale bar, 100 μm. Genotype: *ppk-gal4/+, ppk-cd4-tdtom/uas-ct*^i^. **(B)** Modified Sholl analysis on the morphology of c4da-*wt* and c4da-*ct*^i^. Note that there was a broad region in c4da-*wt* (red bar) in which the dendritic arbor showed a uniform morphology. The shadow areas represented standard deviations. *n*=5 cells for each genotype. The black arrowhead indicated the regions of proximal dendrites. **(C)** The responses of c4da-*ct*^i^ to the force stimuli (4 mN) applied onto the proximal and distal dendrites using a 60 μm probe. The numbers of cells were indicated below the scattered data points. Data were presented as mean±std. *: p<0.05. ns: no significance. *ct*^i^: *ppk-gal4/20×uas-ivs-gcamp6s, ppk-cd4-tdtom/uas-ct*^i^. *scrambled*^i^: *ppk-gal4/20×uas-ivs-gcamp6s, ppk-cd4-tdtom/uas-scrambled*^i^. **(D)** The responses of c4da-*ct*^i^ to the forces applied on distal dendrites using the probes of different sizes. Data were presented as mean±sem (*n*=10 cells). **(E)** The behavioral responses of *ct*^i^ larvae to mechanical poking using the probes of different sizes. *wt*: *w1118*. *ppk-gal4*: *ppk-gal4; +/+*. *uas-ct*^i^: *+/+; uas-ct*^i^. *ct*^i^: *ppk-gal4/+; uas-ct*^i^*/+*. *ppk-gal4* larvae: 30 μm probe (*n*=80 larvae), 60 μm probe (*n*=78 larvae) and 100 μm probe (*n*=75 larvae). *uas-ct*^i^ larvae: 30 μm probe (*n*=81 larvae), 60 μm probe (*n*=75 larvae) and 100 μm probe (*n*=75 larvae). *ct*^i^ larvae: 30 μm probe (*n*=92 larvae), 60 μm probe (*n*=81 larvae) and 100 μm probe (*n*=72 larvae). In panels **C, D** and **E**, the corresponding data from c4da-*wt* were provided for comparison.

As Cut is a transcription factor, the reduction of its expression level might lead to other changes in addition to the disrupted morphology (Jack and DeLotto, 1995; Jinushi-Nakao et al., 2007; Sulkowski et al., 2011). To explore the effects of these potential factors, we performed several control experiments. First, we checked the expression level and subcellular localization of two mechanosensory molecules, i.e. Piezo and Ppk1/Ppk26, in c4da-*ct*^i^. They showed similar expression and localization as in c4da-*wt*. Second, the changes in dendritic cytoskeleton might indirectly affect mechanosensation by altering the mechanical property of dendrites, so we also checked dendritic signals of F-actin and microtubules. No significant differences were found (Fig. s4). These two observations show that although the reduction in the expression level of Cut decreases the number of dendritic branches, the expression and localization of mechanosensory molecules and cytoskeletal elements are unchanged in the existing dendrites of c4da-*ct*^i^. Third, we also performed the same set of mechanical recording experiments on c3da-*wt*, whose mechanosensory machinery is wild-type but dendritic density changes in the way as in the case of c4da-*ct*^i^ (Fig. s2). The responses of c3da-*wt* to the distal stimuli were significantly weaker than those to the proximal stimuli (Fig. s2). In addition, no preference to small probes was found in c3da-*wt* (Fig. s2). These sensory features were markedly different from those in c4da-*wt*, but similar to those in c4da-*ct*^i^. In all, our observations are consistent with the dendritic morphology of c4da making direct contributions in supporting the sensory features. Note that the potential contributions of other unknown factors cannot be absolutely excluded (see Discussion).

### Mechanosensitivity to lateral tension expands the force-receptive field

An unexpected finding in studying c4da-*ct*^i^, primarily due to their sparse dendritic morphology, was that when the force probe did not exert forces on a dendrite but on an adjacent region without any dendrite (i.e. the “dendrite-off” mode, denoted as “***d*-off**” in Fig. 3A), the response of c4da was nearly unchanged, as if the force was directly applied on the dendrite (Fig. 3C). We wondered if this reflects impalpable difference due to the overall reduction in the sensory response of c4da-*ct*^i^ or the dendrites of c4da have an expanded force receptive field. To verify this, we performed the same experiments on c4da-*wt* (Fig. 2D). The responses of c4da-*wt* to the “dendrite-off” stimuli were also unchanged from those to the distal stimuli. Further analysis showed that the responses of c4da kept nearly constant until the force was applied at least 40-60 μm from a dendrite (Fig. 4A). Based on these results, we conclude that the dendrites of c4da have an expanded force-receptive field.

**Figure 4.**
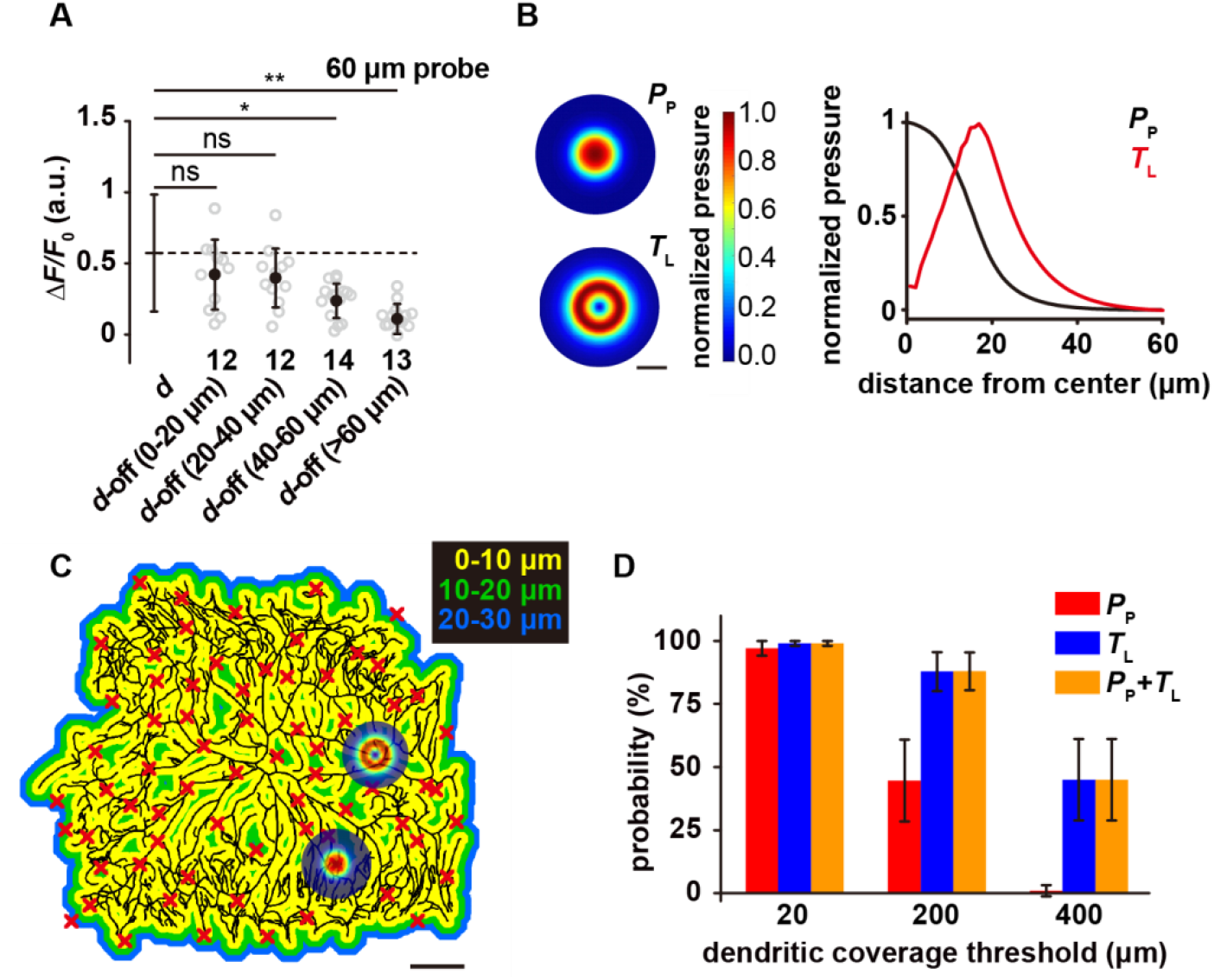
The mechanosensitivity of c4da to lateral tension. **(A)** The responses of c4da to the “dendrite-off” stimuli (4 mN, 60 μm probe) applied at different positions. The numbers of cells were indicated below the scattered data points. Data were presented as mean±std. **: p<0.01. *: p<0.05. ns: no significance. Genotype: *ppk-gal4/+*; *ppk-cd4-tdtom/20×uas-ivs-gcamp6s*. **(B)** Left panel: representative heat maps showing the 2D distributions of pressure perpendicular to the cuticle surface (*P*_P_) (upper) and tension parallel with the cuticle surface (*T*_L_) (lower). Scale bar, 30 μm. Right panel: representative line profiles (normalized) showing the distribution of *P*_P_ and *T*_L_ versus the distance to the center of the force probe. Indentation depth: 20 μm. The diameter of the simulated probe is 60 μm. **(C)** Representative color map for the distance to the nearest dendrite, in which the distance was labeled by different colors as indicated. Random positions were chosen (red cross) as the force application points in our simulations. Scale bar, 100μm. **(D)** The activation probability in three conditions: (1) only sensitive to *P*_P_; (2) only sensitive to *T*_L_; (3) sensitive to both *P*_P_ and *T*_L_. The dendritic coverage threshold (*C*_d_) is the minimal length of activated dendrites that could excite neuronal responses. For each condition, 100 random positions were simulated for each cell. Data were presented as mean±std (*n*=5 cells).

The expansion of force-receptive field suggests that in addition to the sensitivity to the pressure perpendicular to cuticle surface (*P*_p_), c4da also possess mechanosensitivity to lateral tension (parallel to cuticle surface, *T*_L_). By simulating tissue mechanics of larval fillet (Fig. s5, also see Methods), we calculated the distribution of *P*_p_ and *T*_L_ caused by a 60 μm probe (indentation depth: 20 μm, force: ∼ 4 mN). We noted that as the distance to the center of the force probe increased, *P*_p_ showed a monotonic decrease but *T*_L_ peaked at around 20 μm from the force application point (Fig. 4B). This modeling analysis predicts that if c4da were only sensitive to Δ*P*_p_, their sensory response should decrease when the force is not directly on the dendrites. However, if c4da were also sensitive to Δ*T*_L_, the increase in *T*_L_ would compensate for the reduction in *P*_p_. In this scenario, the effective force-receptive field is expanded.

Morphological analysis showed that almost all positions in the dendritic territory of a c4da were within 20 μm distance from a dendrite (Fig. 4C). To examine how the sensitivity to *T*_L_ promotes the likelihood of cell activation, we overlaid the 2D profiles of *P*_p_ and *T*_L_ to the dendrites of c4da at random spots (Fig. 4C) and calculated the probability of cell activation. Note that the calculation had two assumptions. First, we assumed 10% of the peak value of *P*_p_ or *T*_L_ as the activation threshold of a dendritic segment. Second, we assumed a dendritic coverage threshold (*C*_d_), i.e. the minimal amount of excited dendritic segments that would lead to neuronal excitation. To explore the parameter space, we tested three values for *C*_d_, i.e. 0.1% (20 μm) (low threshold), 1% (200 μm) (intermediate threshold) and 2% (400 μm) (high threshold) of the total dendritic length (mean±std: 19560±1667 μm, *n*=5 cells). In simulations, we considered three scenarios, in which the cell is sensitive to either *P*_p_ or *T*_L_ or to both. Simulation results showed that with the lower threshold assumption (e.g. *C*_d_=20 μm), the cells could be activated in all three conditions. However, if the threshold was of intermediate (e.g. *C*_d_=200 μm) or high (e.g. *C*_d_=400 μm) level, the mechanosensitivity to the change in *T*_L_ could largely enhance the probability of neuronal excitation (Fig. 4D), consistent with our hypothesis. In all, our analysis suggests that the sensitivity to lateral tension enhances the efficiency of c4da in force detection, especially when dendritic coverage is small.

### Piezo and Ppk1/Ppk26 differentially contribute to the mechanosensitivity of c4da

We then wondered what is the molecular basis underlying the mechanosensitivity of c4da, especially the sensitivity to lateral tension? Previous behavior assays suggest that Ppk1/Ppk26 and Piezo mediate two parallel mechanosensory pathways in c4da (Kim et al., 2012). This remains to be confirmed at the cellular level. Moreover, it is also unclear how Ppk1/Ppk26 and Piezo contribute to the mechanosensitivity of c4da respectively.

We first confirmed that the dendritic morphologies of c4da-*piezo*^KO^, c4da-*ppk26*^1^ (a null mutant) and c4da-*piezo*^KO^, *ppk1*^Δ5^ (a double null mutant) were not affected (Fig. 5 A-B), excluding the potential effect of morphological changes. We then measured the force-evoked responses of these mutants using two types of probes (30 and 60 μm) and at different dendritic regions (proximal and distal). In all conditions, the responses of c4da-*ppk26*^1^ were significantly reduced and the preference to small probes was completely lost (Fig. 5 C-D). In contrast, the response of c4da-*piezo*^KO^ was changed in a different way. C4da-*piezo*^KO^ showed a mild reduction in the response to the large probe (60 μm) but a much lower response to the small probe (30 μm). This phenotype was more prominent for the distal stimuli than for the proximal stimuli (Fig. 5 C-D). As a result, the preference to small probes was only attenuated for the proximal stimuli but completely lost for the distal stimuli (Fig. 5 C-D). Based on these measurements, we conclude that while Ppk1/Ppk26 contributes to the overall mechanosensitivity of c4da, Piezo is particularly important for detecting more localized forces. Furthermore, c4da-*piezo*^KO^, *ppk1*^Δ5^ (the double mutant) showed almost no response to any force stimuli (Fig. 5E), suggesting that the contributions of Ppk1/Ppk26 and Piezo are additive. Finally, the responses of c4da-*piezo*^KO^ and c4da-*ppk26*^1^ at the “dendrite-off” mode were comparable to the responses to the distal stimuli, showing that both mutants are still able to sense lateral tension (Fig. 5F). However, their responses to the “dendrite-off” stimuli were both significantly weaker than that of c4da-*wt* (Fig. 5F), suggesting that Ppk1/Ppk26 and Piezo both make contributions to the lateral mechanosensitivity of c4da.

**Figure 5.**
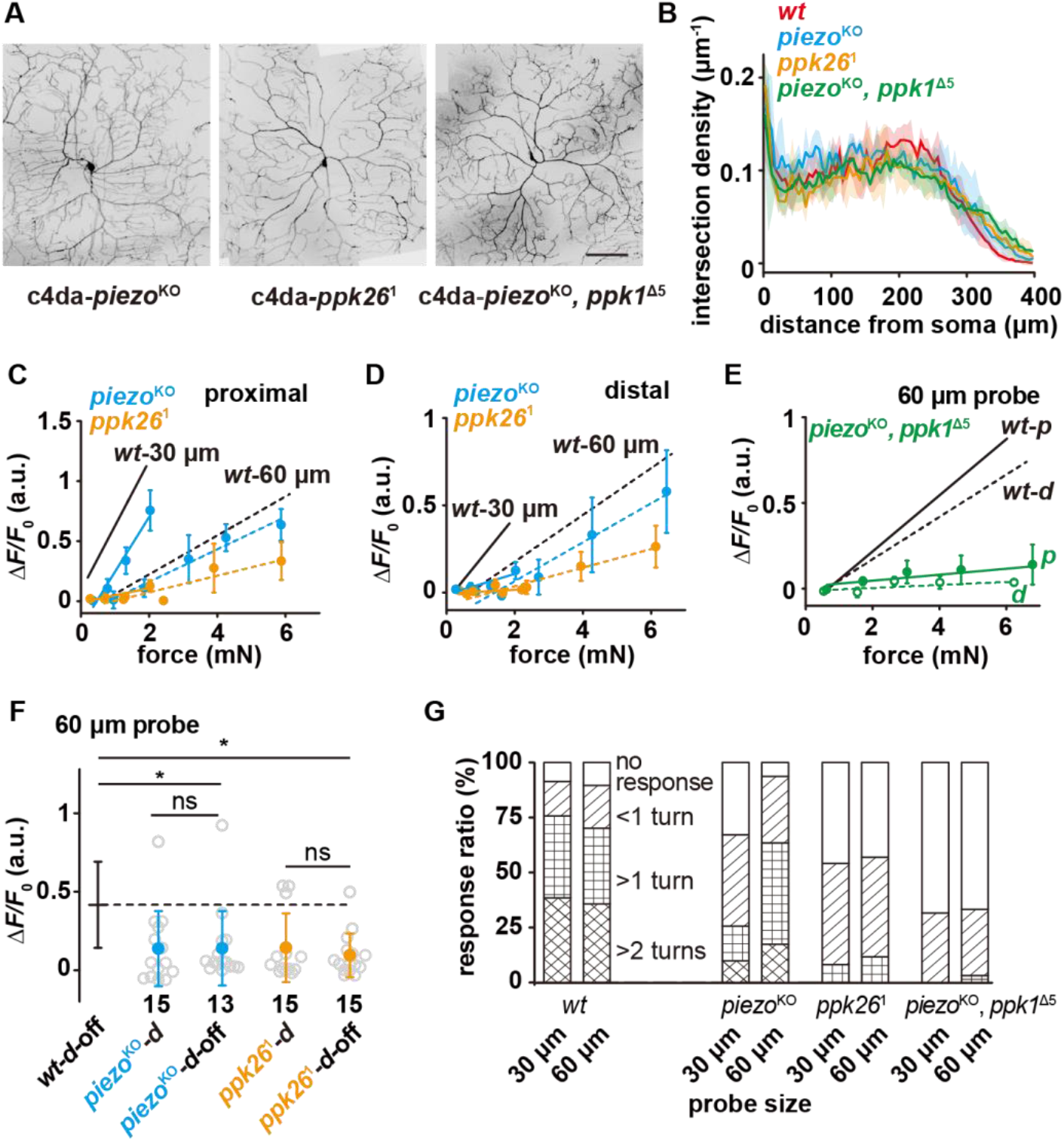
The differential contributions of Piezo and Ppk1/Ppk26 to the mechano-sensitivity of c4da. **(A)** Representative images of c4da-*piezo*^KO^ (*piezo*^KO^; *ppk-cd4-tdtom/+*), c4da-*ppk26*^1^ (*uas-cd4-tdtom/ppk-gal4*; *ppk26*^1^), c4da-*piezo*^KO^*, ppk*^Δ5^ (*piezo^KO^*, *ppk*^Δ5^; *ppk-cd4-tdtom/+*). Scale bar: 100 μm. **(B)** Modified Sholl analysis on the morphologies of c4da-*wt* (*n*=5), c4da-*piezo*^KO^ (*n*=3), c4da-*ppk26*^1^ (*n*=3) and c4da-*piezo*^KO^, *ppk*^Δ5^ (*n*=3). The shadow areas represented standard deviations. **(C)** and **(D)** The responses of c4da-*piezo*^KO^ and c4da-*ppk26*^1^ to the proximal (**C**) and distal **(D)** stimuli delivered using the probes of two different sizes. Solid lines: 30 μm probe. Dashed lines: 60 μm probe. *piezo*^KO^: *piezo*^KO^*; ppk-gal4/20×uas-ivs-gcamp6s* (*n*=12 cells). *ppk26*^1^: *ppk-gal4/20×uas-ivs-gcamp6s; ppk26*^1^ (*n*=12 cells). **(E)** The responses of c4da-*piezo^KO^*, *ppk*^Δ5^ to the proximal and distal stimuli delivered using a 60 μm probe. Solid lines: proximal stimuli. Dashed lines: distal stimuli. Genotype: *piezo*^KO^, *ppk1*^Δ5^: *piezo*^KO^, *ppk1*^Δ5^*; ppk-gal4/20×uas-ivs-gcamp6s* (*n*=8 cells). **(F)** The responses of c4da-*piezo*^KO^ and c4da-*ppk26*^1^ to the “dendrite-off” stimuli (4 mN) delivered using a 60 μm probe. The numbers of cells were indicated below the scattered data points. Data were presented as mean±std. *: p<0.05. ns: no significance. **(G)** The behavioral responses of the c4da-*piezo*^KO^, c4da-*ppk26*^1^ and c4da-*piezo*^KO^, *ppk*^Δ5^ larvae to mechanical poking using the probes of different sizes. *wt*: *w1118*. *piezo*^KO^: *piezo*^KO^*; +/+*. *ppk26*^1^: *+/+; ppk26*^1^. *piezo*^KO^, *ppk1*^Δ5^: *piezo*^KO^, *ppk1*^Δ5^*; +/+*. *piezo*^KO^ larvae: 30 μm probe (*n* = 77 larvae), 60 μm probe (*n* = 78 larvae). *ppk26*^1^ larvae: 30 μm probe (*n* = 77 larvae), 60 μm probe (*n* = 74 larvae). *piezo*^KO^, *ppk1*^Δ5^ larvae: 30 μm probe (*n* = 62 larvae), 60 μm probe (*n* = 63 larvae). In panel **C**, **D** and **E,** data were presented as mean±sem. In panels **C, D**, **E, F** and **G**, the corresponding data from c4da-*wt* were provided for comparison.

To further verify the physiological contributions of Ppk1/Ppk26 and Piezo in mechanical nociception, we performed the behavior assay of mechanical nociception on these three mutants using the 30 and 60 μm probes. C4da-*piezo*^KO^ showed a largely reduced response to the 30 μm probe but an almost unchanged response to the 60 μm probe. Meanwhile, c4da-*ppk26*^1^ showed very weak responses to both of the probes and the double mutant (i.e. c4da-*piezo*^KO^, *ppk1*^Δ5^) showed almost no responses to all mechanical stimuli (Fig. 5G). In all, these behavioral observations are consistent with our cellular observations.

### Ca-α1D contributes to the dendritic signal propagation in c4da

Finally, we addressed if the cellular mechanosensitivity of c4da relies on active signal propagation in the dendrites. The unchanged responses to the distal and proximal stimuli using the 60 μm probe (Fig. 2E) support the idea of active propagation in the dendrites. Meanwhile, the lower response to the distal stimuli than that to the proximal stimuli using the 30 μm probe (smaller contact interface) in both c4da-*wt* (Fig. 2E) and c4da-*piezo*^KO^ (Fig. 5 C-D) suggests that the dendritic signal propagation mechanism is not all-or-none but graded, likely dependent on the amount of dendritic segments being activated. This hypothesis predicts that if the dendritic signal propagation is weakened, the cellular response of c4da to smaller probes would be affected to a greater extent. We then tested this hypothesis.

A previous study showed that Ca-α1D, a voltage-gated calcium channel (VGCC), contributes to the heat-evoked calcium transient in the dendrites of c4da (Terada et al., 2016), suggesting its function in facilitating dendritic signal propagation. Therefore, we first examined the roles of VGCCs in force-evoked responses. We noted that upon the mechanical stimuli delivered using a 60 μm probe, calcium increase can be observed in the dendrites around the force application point and the soma (Fig. 6A), reflecting the propagating signals in the dendrites. The response in the dendrites can be suppressed by nimodipine (5 μM), an antagonist of VGCCs, or by knocking down the expression level of *Ca-α1D* (*Ca-α1D*^i^) (Fig. 6 A-B). Therefore, Ca-α1D also contributes to dendritic signal propagation in the force-evoked responses of c4da.

**Figure 6.**
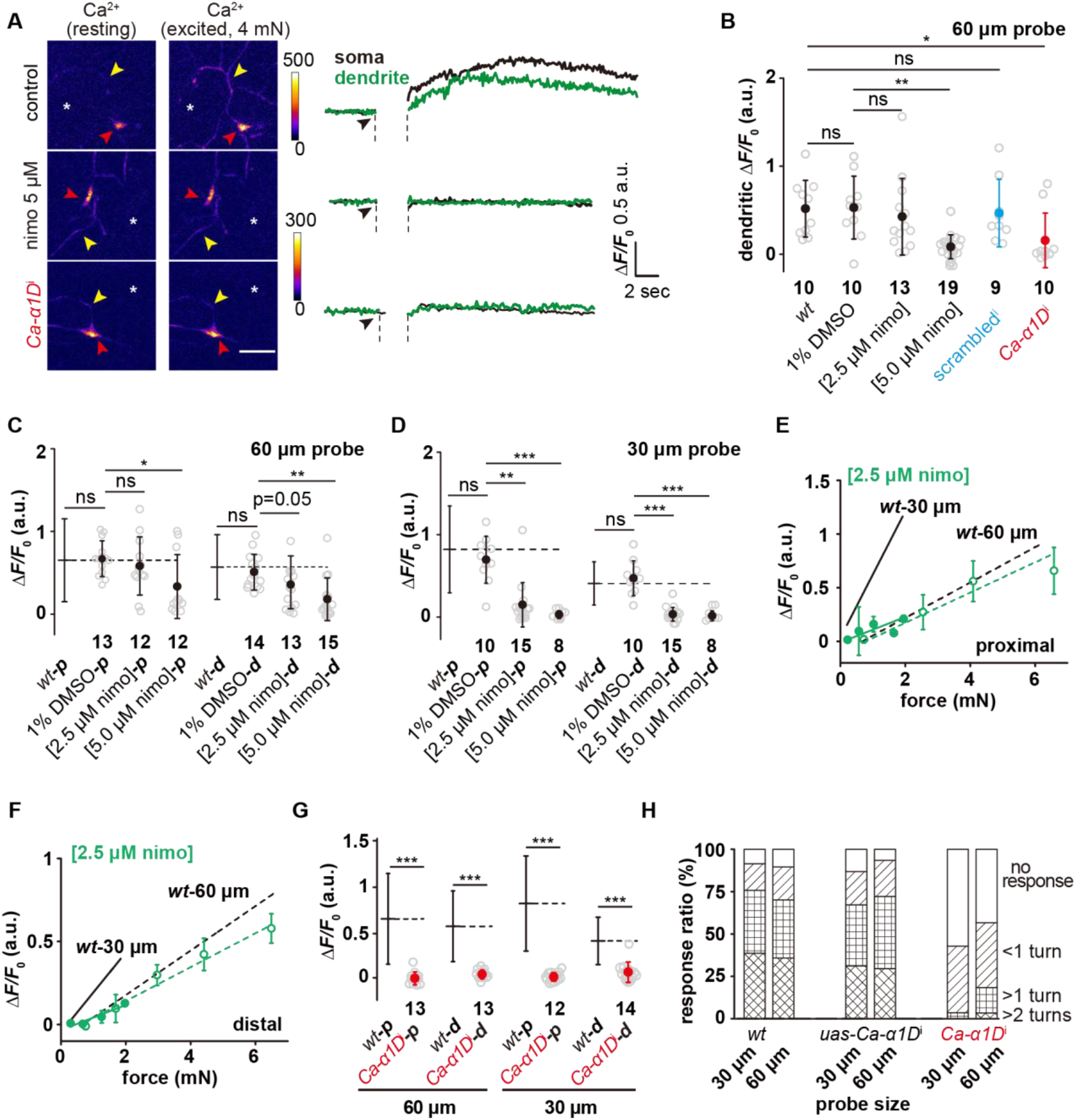
Active signal propagation in the dendrites of c4da. **(A)** Representative images and curves showing the somatic (red arrowhead) and dendritic (yellow arrowhead) responses (calcium increase) of c4da to the stimuli (4 mN, 60 μm probe) in c4da-*wt*, nimodipine-treated c4da-*wt* and c4da-*Ca-α1D*^i^. The asterisk indicated the force application point. Scale bar: 50 μm. C4da-*wt* and nimodipine treated: *ppk-gal4/+*; *ppk-cd4-tdtom/20×uas-ivs-gcamp6s*. *Ca-α1D*^i^: *ppk-gal4/20×uas-ivs-gcamp6s*; *ppk-cd4-tdtom/ uas-Ca-α1D*^i^. **(B)** Statistics quantification of calcium increase in the dendrites in response to the stimuli (4 mN, 60 μm probe). C4da-*wt* and nimodipine treated c4da-*wt*: *ppk-gal4/+*; *ppk-cd4-tdtom/20×uas-ivs-gcamp6s*. *Ca-α1D*^i^: *ppk-gal4/20×uas-ivs-gcamp6s*; *ppk-cd4-tdtom/uas-Ca-α1D*^i^. *Scrambled*^i^: *ppk-gal4/20×uas-ivs-gcamp6s, ppk-cd4-tdtom/uas-Scrambled*^i^. **(C)** and **(D)** The responses of nimodipine treated c4da-*wt* to the proximal and distal stimuli to the probes of different sizes. **(C)**: 60 μm probe stimuli. **(D)**: 30 μm probe stimuli. **(E)** and **(F)** The responses of c4da-*wt* (*n*=10 cells) treated with 2.5 μM nimodipine to the proximal (**E**) and distal **(F)** stimuli delivered using the probes of two different sizes. Solid lines: 30 μm probe. Dashed lines: 60 μm probe. **(G)** The responses of c4da-*Ca-α1D*^i^ to the stimuli applied at different dendritic regions (proximal and distal) and delivered using the probes of two different sizes (60 μm: 4 mN, 30 μm: 1.5 mN). **(H)** The behavioral responses of the c4da-*Ca-α1D*^i^ larvae to mechanical poking using the probes of two different sizes. *wt*: *w1118*. *ppk-gal4*: *ppk-gal4; +/+*. *uas-Ca-α1D*^i^: *+/+; uas-Ca-α1D*^i^*. Ca-α1D*^i^: *ppk-gal4/+;uas-Ca-α1D*^i^ */+*. *uas-ct*^i^ larvae: 30 μm probe (*n*=72 larvae), 60 μm probe (*n*=71 larvae). *ct*^i^ larvae: 30 μm probe (*n*=65 larvae), 60 μm probe (*n*=71 larvae). In panels **B**, **C**, **D**, **G**, the numbers of cells were indicated below the scattered data points and data were presented as mean±std. ***: p<0.001. **: p<0.01. *: p<0.05. ns: no significance. In panel **E** and **F,** data were presented as mean±sem. In panels **C, D**, **E**, **F, G** and **H**, the corresponding data from c4da-*wt* were provided for comparison.

Drug titration experiments showed that in the condition of using a 60 μm probe, 5 μM nimodipine, the full inhibition concentration in the literatures (Scriabine and van den Kerckhoff, 1988; Terada et al., 2016), significantly suppressed the responses of c4da to both proximal and distal stimuli, while 2.5 μM nimodipine showed no effect on the response to the proximal stimuli but a moderate suppression on that to the distal stimuli (Fig. 6C). We interpreted this observation as when the VGCCs in the dendrites of c4da were partially inhibited, the responses to the distal stimuli were affected to a greater extent, in consistence with the idea that active signal propagation is more important in sensing distal stimuli. When a 30 μm probe was used, 2.5 μM nimodipine showed an inhibitory effect on the response to the proximal stimuli and an even stronger inhibition on that to the distal stimuli (Fig. 6D). Further measurements on the responses of c4da to the full range of stimulating forces in the presence of 2.5 μM nimodipine showed that the drug largely suppressed the responses to the 30 μm probe but had only a mild effect on those to the 60 μm probe (Fig. 6 E-F). In this condition, the sensory preference of c4da to small probes was completely lost (Fig. 6 E-F), consistent with the idea that the inhibition of the VGCCs reduces the responses to small probes to a greater extent.

In comparison to c4da-*wt*, c4da-*Ca-α1D*^i^ showed significantly reduced responses to the stimuli delivered using the probes of two different sizes (30 and 60 μm) or at different dendritic regions (Fig. 6G), suggesting that Ca-α1D is likely a major contributing VGCC for active signal propagation in the dendrites of c4da. Using the behavior assays of mechanical nociception, we found that the defensive response of the *Ca-α1D*^i^ larvae in response to mechanical poking was largely weakened, especially the responses to the smaller probes (Fig. 6H). These results accord with our cellular observations.

## Discussion

In the present study, we show that c4da are sensitive to mN-scale forces and uniformly respond to the forces applied at different dendritic regions. In particular, c4da appear to be more sensitive to small probes, given that the applied forces are the same. We reveal three cellular mechanisms that facilitate the sensory features of c4da (Fig. 7). First, the high morphological complexity ensures dense and uniform distribution of mechanosensory molecules across the entire dendritic tree. Second, the mechanosensitivity to lateral tension, which depends on Ppk1/Ppk26 and Piezo, expands the force-receptive field of the dendrites. Third, the active signal propagation in the dendrites promotes the overall mechanosensitivity of c4da and has a more prominent effect on the sensitivity to small probes. Ca-α1D, a voltage-gated calcium channel, is a major contributing molecule in mediating the dendritic signal propagation. We now discuss the potential implications of these findings in understanding the mechanosensory and nociceptive functions of c4da.

**Figure 7.**
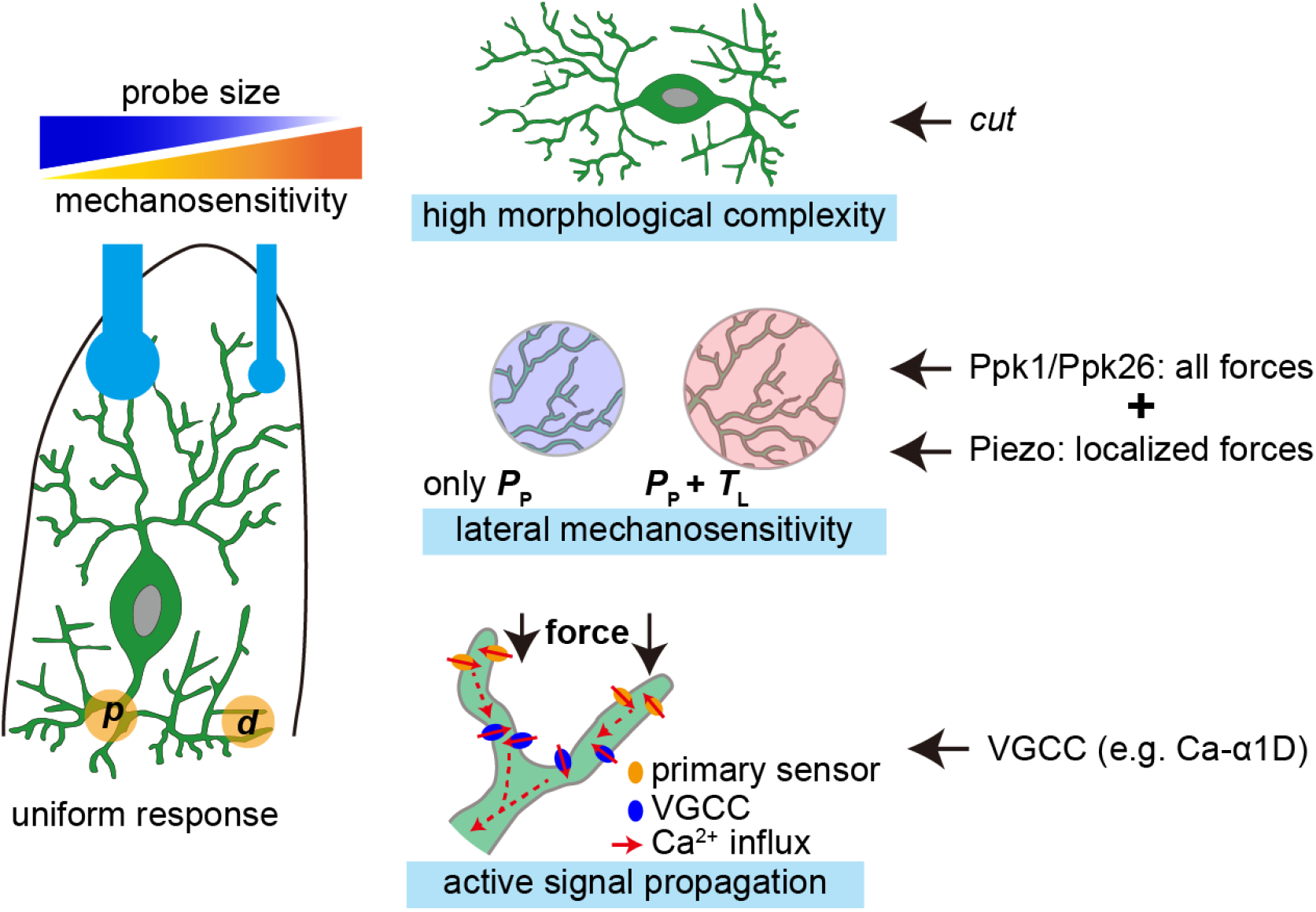
The mechanisms underlying the mechanosensory features of c4da. Left panel: c4da showed a greater sensitivity to localized poking forces and made uniform responses to the forces applied at different dendritic regions (***p***: proximal stimuli. ***d***: distal stimuli). Right panel: the key cellular mechanisms that facilitate the sensory features of c4da and the important contributing molecules.

### Implications for the mechano-nociceptive functions of c4da

C4da are first implicated as mechanoreceptor cells using the behavior assays (Gorczyca et al., 2014; Guo et al., 2014; Kim et al., 2012; Mauthner et al., 2014; Zhong et al., 2010). Cellular responses to force stimuli are observed on isolated *ppk1*-positive cells and whole mount larval fillet preparations (Guo et al., 2014; Kim et al., 2012; Tsubouchi et al., 2012; Walcott et al., 2018; Yan et al., 2013). Here, we provide quantitative characterizations on the responses of c4da to the force stimuli with controlled strength, contact area and force application position. Our conclusions add to the current model of understanding how c4da act as a mechanical nociceptor in two aspects.

At the cellular level, we find that with the same force, c4da show a greater sensitivity to smaller probes than to larger probes. This provides a cellular basis to understand the physiological function of c4da in sensing mechanical poking from sharp objects, e.g. the ovipositor of wasp (Hwang et al., 2007). Geometry of the force probes determines that for a given change of force, smaller probes caused a smaller Δ*A*_c_ but a greater Δ*P* in comparison to larger probes. We find that c4da develop several cellular mechanisms to ensure that the pressure change over a small contact interface could be collected, amplified and finally encoded into neuronal responses. First, the high dendritic density allows an adequate dendritic coverage for small probes. Second, the lateral mechanosensitivity expands the force-receptive field of dendrites, acting as a mechanical amplifier in receiving a localized force. Third, the active signal propagation, as an intracellular electrochemical amplifier, compensates for signal attenuation in the dendrites. These mechanisms optimize the mechanosensitivity of c4da from structural, mechanical and signaling aspects, respectively. In essence, they all serve to increase the probability of activating the mechanosensory pathways that are required to excite neuronal responses.

At the molecular level, we find that Piezo and Ppk1/Ppk26 make differential contributions to the mechanosensitivity of c4da. Previous behavior assays suggest that Ppk1/Ppk26 and Piezo in c4da mediate two parallel pathways in mechanical nociception (Gorczyca et al., 2014; Guo et al., 2014; Kim et al., 2012; Mauthner et al., 2014). This raises the question of why c4da need two sets of mechanosensory pathways. Furthermore, patch clamp recordings show that the force-evoked electrical response of isolated *ppk*-positive cells is entirely dependent on Piezo (Kim et al., 2012), which contradicts the inference based on the behavior assays. Due to the unsolved issues and inconsistency, it is necessary to explore how Ppk1/Ppk26 and Piezo contribute to the in-vivo mechanosensitivity of c4da. Here, we discover that Piezo is particularly important for c4da to detect localized forces but plays a relatively minor role in sensing large probes. In contrast, the loss of Ppk1/Ppk26 reduces the responses of c4da to the entire range of stimulating forces used in our experiments, suggesting the contribution of Ppk1/Ppk26 to the overall mechanosensitivity. In addition, the lack of responses in the double knock-out mutant strain confirms that the contributions of Ppk1/Ppk26 and Piezo are additive, as suggested by the previous behavior assays. Therefore, our findings add to the current model by showing a more specific role of Piezo. Furthermore, we also show that Ppk1/Ppk26 and Piezo both contribute to the sensitivity of c4da to lateral tension. In consistence with this idea, members in the Piezo and the DEG/ENaC channel families have been proposed to be gated by the changes in lateral membrane tension (Cueva et al., 2007; Guo and MacKinnon, 2017; Liang and Howard, 2018; Lin et al., 2019). However, it remains unclear how Piezo could facilitate the greater sensitivity to localized forces. The underlying mechanism may reside in the gating property of Piezo, such as gating sensitivity and conductance. This issue awaits future studies.

### Contribution of dendritic morphology

Based on the observations in the *ct*^i^ strain (Fig. 3), we argue that the morphological complexity is a key factor in supporting the sensory features of c4da. We note that Cut is a transcription factor that contributes to the morphological determination of c4da. Is it possible that the altered mechanosensory responses observed in the *ct*^i^ mutant is due to other downstream changes? While this possibility cannot be absolutely excluded, we think that the contribution of dendritic morphology is valid based on several lines of evidence and thoughts. First, the responses of c4da-*ct*^i^ to the stimuli on the proximal dendritic region, where the dendritic density is similar to that of c4da-*wt*, is comparable to those of c4da-*wt*, suggesting that the essential mechanosensory elements are fairly normal in the existing dendrites of c4da-*ct*^i^ (Fig. 3C). Second, the expression and localization of mechanosensory molecules (Piezo and Ppk1/Ppk26) and cytoskeletal elements (F-actin and microtubule) in c4da-*ct*^i^ are comparable to those in c4da-*wt* (Fig. s4), although it is formally possible that the fine organization of these molecular components could still be different in c4da-*ct*^i^. Third, at least in terms of how a cell responds to different probe sizes and to the stimuli at different dendritic regions, c4da-*ct*^i^ are similar to c3da-*wt*, whose morphological pattern is markedly different from c4da-*wt* but close to c4da-*ct*^i^ (Fig. s2). This similarity, if neglecting the differences between c3da and c4da at the molecular level, reflects the contribution of dendritic morphology.

## Materials and Methods

### Fly stocks

All flies were maintained on standard medium at 23-25℃ in 12:12 light-dark cycles. *20×uas-ivs-gcamp6s* (BL42746, BL42749), *ppk-gal4* (BL32078, BL32079), *uas-cd4-tdTom* (BL35837, BL35841), *ppk-cd4-tdtom* (BL35845), *ppk-cd4-tdgfp* (BL35843) and *uas-lifeact-rfp* (BL58715) were from Bloomington Drosophila Stock Center (BDSC). *Gal4-19-12* was from the Jan Lab (UCSF). *ppk26-gfp* was from the Tracey Lab (Indiana University). *Uas-mcherry-jupiter* was from the Han Lab (Cornell University). *Tmc-gal4*, *piezo-gfp*, *piezo*^KO^, *ppk26*^1^ and *piezo*^KO^*, ppk*^Δ5^ and were from Wei Zhang (Tsinghua University). *Ct*^i^ (THU1309), *Ca-α1D*^i^ (THU0766) and the control strains were from Tsinghua Fly Center.

### Larval fillet preparation

Third instar larvae were dissected on a 35 mm dish coated with a PDMS pad (thickness: 1 mm) in the insect hemolymph-like (HL) buffer (Stewart et al., 1994). The cuticle, epidermis and muscle tissue were kept as intact as possible. The fillet was mounted on the PDMS pad using 8-10 insect pins (Austerlitz Insect Pins, Czech) and kept as stretched. HL buffer: 103 mM NaCl (10019318 Sinopharm Chemical Reagent Co., Ltd., Shanghai, China), 3 mM KCl (P9541 Sigma-Aldrich, USA), 5 mM TES (T5691 Sigma-Aldrich, USA), 8 mM trehalose (T1067 Sigma-Aldrich, USA), 10 mM glucose (G7528 Sigma-Aldrich, USA), 5 mM sucrose (V900116 Sigma-Aldrich, USA), 26 mM NaHCO_3_ (A500873 Sangon Biotech, Co., Lt., Shanghai, China), 1 mM NaH_2_PO_4_ (S8282 Sigma-Aldrich, USA), 2 mM CaCl_2_ (Xilong Scientific Co., Ltd., Shantou, China), and 4 mM MgCl_2_ (M8266 Sigma-Aldrich, USA). The PDMS pad was made using the Sylgard 184 kit (Dow, UAS).

### Mechanical device setup and calibration

The mechanical device consisted of a piezo actuator (PZT 150/7/60 VS12, SuZhou Micro Automation Technology Co., Ltd., Suzhou, China), a strain gauge (customized in Nanjing Bio-inspired-tech Co., Ltd., Nanjing, China) and a force probe. The stepping distance (*D*) of the piezo actuator and the deflection of the metal beam (*B*) were measured using a digital camera with a pixel size of 1.1 μm (AO-3M630, AOSVI optical instrument Co., Ltd. Shenzhen, China). The voltage readout of the strain gauge after the addition of pure water with a minimal step of 100 μL (i.e. 0.98 mN) was recorded using a data acquisition card (NBIT-DSU-2404A, Nanjing Bio-inspired-tech Co., Ltd., Nanjing, China). The force probes were made from capillary glass tubes using an electrode puller (PC-10, Narishige, Japan) and then polished using a microforge (MF-830, Narishige, Japan). Finally, the assembled mechanical device was mounted on the working stage of the spinning-disk confocal microscope (Andor, UK).

### Confocal microscopy

The calcium fluorescent signals were recorded using an Andor inverted spinning disk confocal system (Andor, UK) equipped with an inverted microscope (Olympus, IX73), an iXon 897 EMCCD and a long working-distance 20× objective (UCPlanFL N, N.A. 0.70, working distance: 2.1 mm) (Olympus, Japan).

The images of Piezo-GFP, Ppk26-GFP, mCherry-Jupiter and Lifeact-RFP were acquired using a Zeiss 780 confocal microscope equipped with a 63× objective (Zeiss Plan-Apochromat, 1.4 N.A., Germany) and GaAsP detectors (Zeiss, Germany).

### Scanning electron microscopy

The scanning electron microscopy (SEM) pictures of the glass probes were imaged using FEI Quanta 200 (Thermofisher, USA) with 15-kV voltages and 500×magnification in the high vacuum mode.

### Modified behavior assay

The Von Frey fiber (6lb. Omniflex Line, Zebco) was mounted onto a holder. The free fiber was 20 mm long. The customized glass probe was then fixed at the free end of the fiber with ergo 5400 (Kisling, Switzerland). The poking forces were measured using an electronic balance (XPR204S, Mettler Toledo, USA). The behavior assays were carried on as the previously described (Hwang et al., 2007; Zhong et al., 2010). Briefly, we used the modified fiber to poke the dorsal side of larval middle segments and recorded their behavioral responses, which were then graded for statistical analysis (rolling over 2 turns, rolling over 1 turn, rolling less than 1 turn or no response).

### Drug treatment

Nimodipine (N149 Sigma-Aldrich, USA) was dissolved in DMSO (276855 Sigma-Aldrich, USA). The incubation time for all drugs was 20 min.

### The “sphere-surface” contact mechanics model

Based on the classical “sphere-surface” contact model (Johnson, 1985), we had

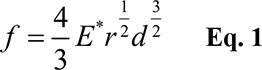

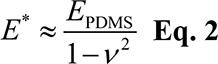

where *f* was the total force, *r* was the radius of the sphere, *E*_PDMS_ was the elastic modulus of PDMS, *E*\* was the effective modulus, *ν* was the Poisson’s ratio of PDMS and *d* was the indentation depth. *d* can be calculated as

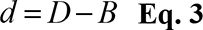

where *D* was the stepping displacement generated by the piezo stack actuator and *B* was the bending deflection of the metal beam (Fig. 1A). In this model, the pressure at the center of contact area (*P*_0_) can be calculated as

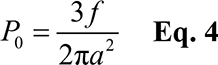

where *a* was the radius of the contact area and could be calculated as

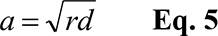

The area of contact surface (*A*_c_) can be calculated as

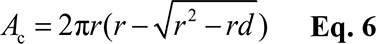

The pressures at a given point to the soma can be calculated as

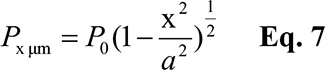

where *x* (μm) was the distance to the center of contact area.

### Finite element analysis

To theoretically investigate the stress field in the cuticle caused by indentation, finite element simulations were performed using the commercial software ABAQUS 6.14.1. (Dassault Systèmes, France). Due to symmetry of the load and the geometry, an axisymmetric mechanical model was established (Fig. s5). The model includes the spherical probe and the underlying composite that consists of two layers of materials: (1) i.e. the cuticle layer (thickness 10 μm, elastic modulus *E*_cuticle_=10 MPa, Poisson’s ratio ν_cuticle_=0.45); (2) the underlying PDMS substrate (thickness 1000 μm, elastic modulus *E*_PDMS_=2.6 MPa, Poisson’s ratio ν_PDMS_=0.45). Because the muscle layer (elastic modulus *E*_muscle_=10 kPa) was much more compliant than cuticle and PDMS (Kot et al., 2012), it is expected to make little contribution to the force distribution. Therefore, we omitted the muscle layer in our finite element model for simplicity. The probe was treated as a rigid object, while the cuticle layer and PDMS substrate were assumed to be linearly elastic. Displacement load in the vertical direction was applied to the probe. The lower surface of the cuticle layer and the upper surface of the PDMS substrate were assumed to be tightly coupled. Fixed boundary conditions were applied to the bottom surface of the PDMS substrate. The compressive pressure (perpendicular to cuticle surface) and lateral (parallel to cuticle surface) tension were both calculated. The simulation of random positioned stimuli was performed using MATLAB (MathWorks, USA).

### Image analysis

GCaMP6s signal was measured using Fiji (Schindelin et al., 2012). The calibration bars and scale bars were generated in Fiji. Sholl analysis on neuronal morphology was carried out using Fiji (Ferreira et al., 2014). The density of intersections was calculated as the number of intersections divided by the circumference of corresponding circle in the Sholl analysis (Fig. 3, Fig. 5 and Fig. s2).

### Conventional data plotting and statistical analysis

Data plotting and statistical analysis were performed using Origin (OriginLab, UAS). The heat maps of the *P*_P_ and *T*_L_ were generated using MATLAB. Statistical analysis (Student’s *t* test) was performed using Origin.

## Supplementary Materials

**Figure s1.**
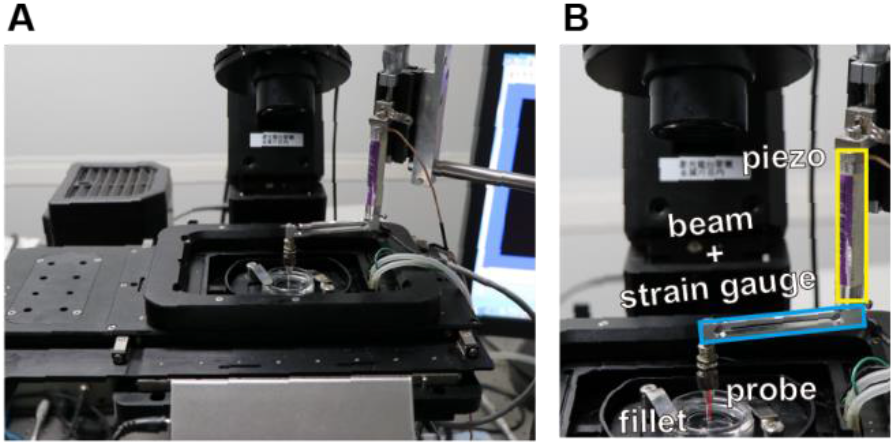
The customized mechanical device. **(A)** The mechanical device was mounted on the working stage of a spinning-disk confocal microscope. **(B)** The mechanical device. Yellow box: the piezo actuator. Cyan box: the beam coupled with a strain gauge. Red bar: the glass probe.

**Figure s2.**
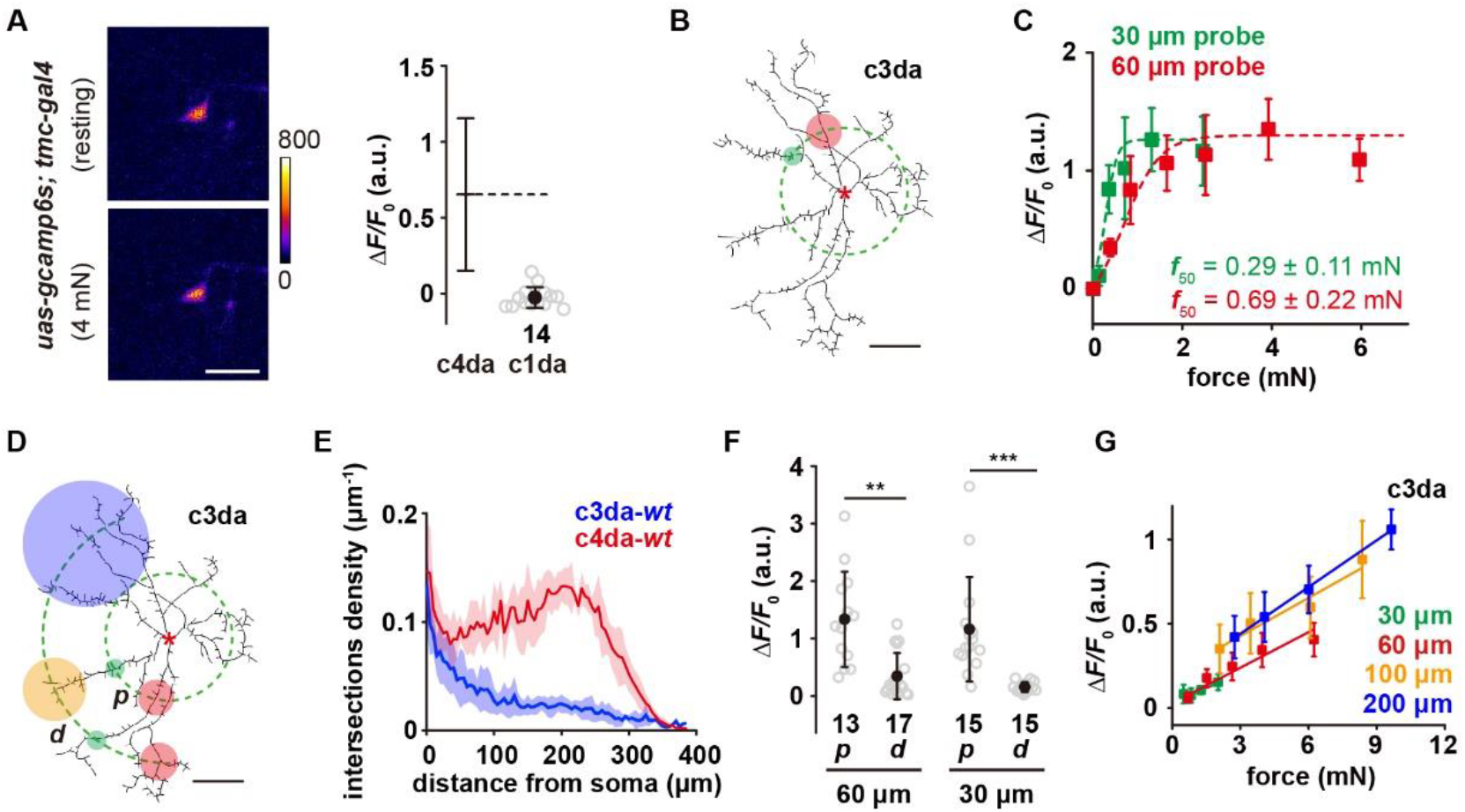
Mechanosensory responses of c1da and c3da. **(A)** Left panel: the representative image showing that c1da make no response to a 4 mN force stimulus. Right panel: Statistical quantification of the responses of c1da to compressive forces (4 mN, 60 μm probe). **(B)** A representative image of c3da. The forces were applied at about 100 μm from the soma, i.e. along the green dashed circle. The representative force application points were marked using the filled circles (Green: 30 μm probe. Red: 60 μm probe). The data of c4da-*wt* (see Fig. 2E) were provided for comparison. Genotype: *uas-cd4-tdtom; gal4-19-12*. **(C)** The force-response (Δ*F*/*F*_0_) plots of c3da (*n*=12 cells) using the probes of two different sizes (Green: 30 μm probe. Red: 60 μm probe). The dashed lines were Boltzmann fitting. **(D)** The schematic showing the stimuli (filled circles, green: 30 μm probe, red: 60 μm probe, orange: 100 μm probe, blue: 200 μm probe) delivered at different positions using the probes of different sizes on c3da. ***p***: proximal dendrite. ***d***: distal dendrite. **(E)** Modified Sholl analysis on the morphology of c3da-*wt* and c4da-*wt*. The shadow areas represented standard deviations. *n*=5 cells for each type of neurons. **(F)** The responses of c3da to the proximal and distal stimuli using the 30 (1.5 mN) and 60 μm (4 mN) probes. **(G)** The responses of c3da (*n*=9 cells) to the forces applied on distal dendrites using the probes of different sizes. In panel **A**, scale bar: 30 μm. In panel **A** and **F**, data were presented as mean±std and the numbers of cells were indicated below the scattered data points. In panel **B**, and **D**, scale bar: 100 μm. In panel **C** and **G**, data were presented as mean±sem. Asterisk: the soma. ***: p<0.001. **: p<0.01.

**Figure s3.**
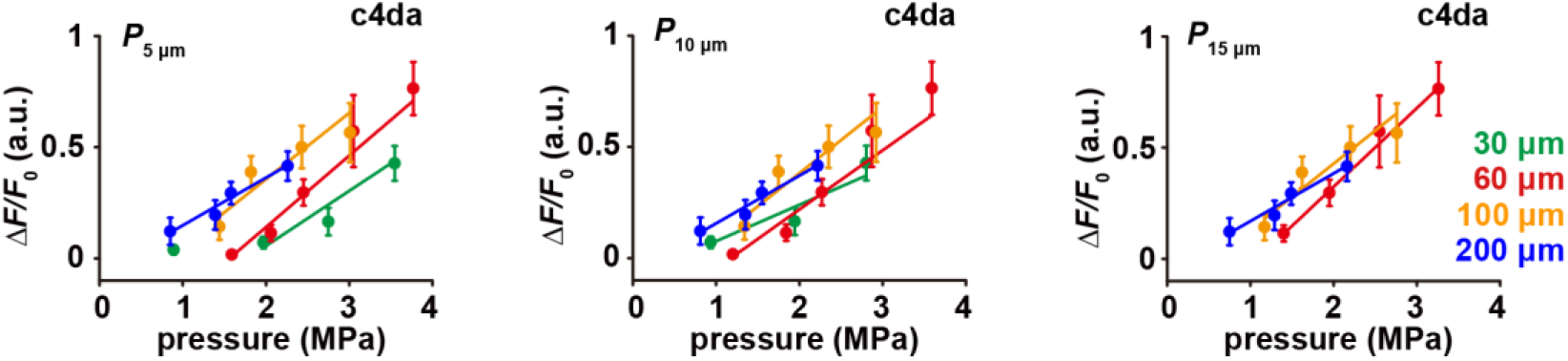
The responses of c4da-*wt* to distal stimuli have a linear scaling relationship with the pressure (*P*_x μm_) Left panel: The responses of c4da to distal stimuli versus the pressures measured at 5 μm, 10 μm and 15 μm away from the center of the force (*P*_5 μm_, *P*_10 μm_, *P*_15 μm_). Data were presented as mean±sem (*n*=10 cells).

**Figure s4.**
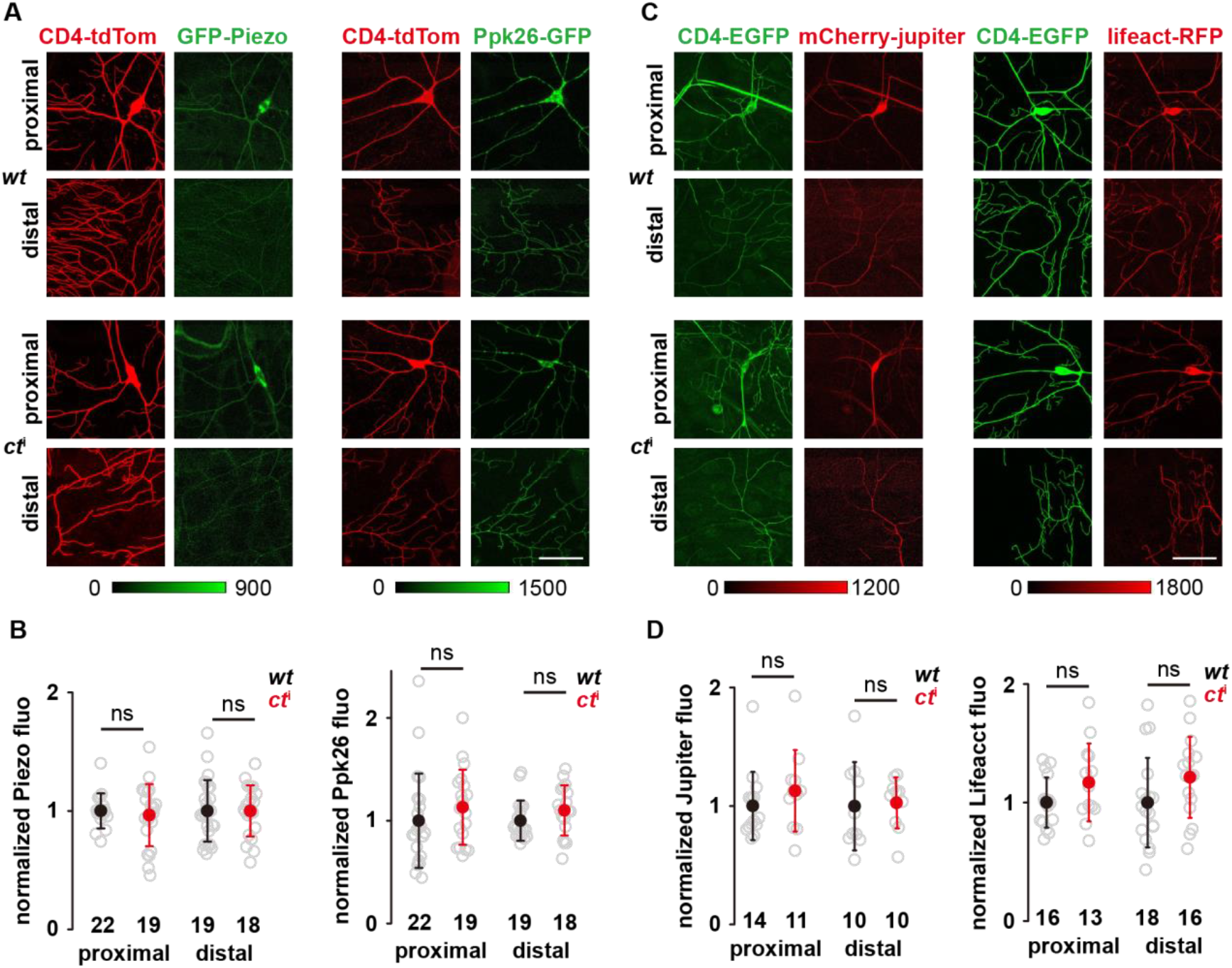
The expression and localization of the mechanosensory molecules and cytoskeletal elements in c4da-*ct*^i^. **(A)** The representative images of the mechanosensory molecules in c4da-*wt* and c4da-*ct*^i^. G*fp-piezo*: *gfp-piezo/+; ppk-cd4-tdtom/+*. *Ppk26-gfp*: *ppk-gal4*/+; *ppk-cd4-tdtom/uas-ppk26-gfp*. **(B)** Statistical quantification of Piezo-GFP (left panel) and Ppk26-GFP (right panel) signals in c4da-*wt* and c4da-*ct*^i^. **(C)** The representative images of f-actin (lifeact) and microtubules (jupiter) in c4da-*wt* and c4da-*ct*^i^. *Mcherry-jupiter*: *ppk-gal4/+; ppk-cd4-tdgfp/uas-mcherry-jupiter*. *Lifeact-rfp*: *ppk-gal4*/+; *ppk-cd4-tdgfp /uas-lifeact-rfp*. **(D)** Statistical quantification of mCherry-Jupiter (left panel) and Lifeact-RFP (right panel) signals in c4da-*wt* and c4da-*ct*^i^. In panel **A** and **C**, scale bar: 50 μm. In panel **B** and **D**, data were presented as mean±std and the numbers of cells were indicated below the scattered data points. ns: no significance.

**Figure s5.**
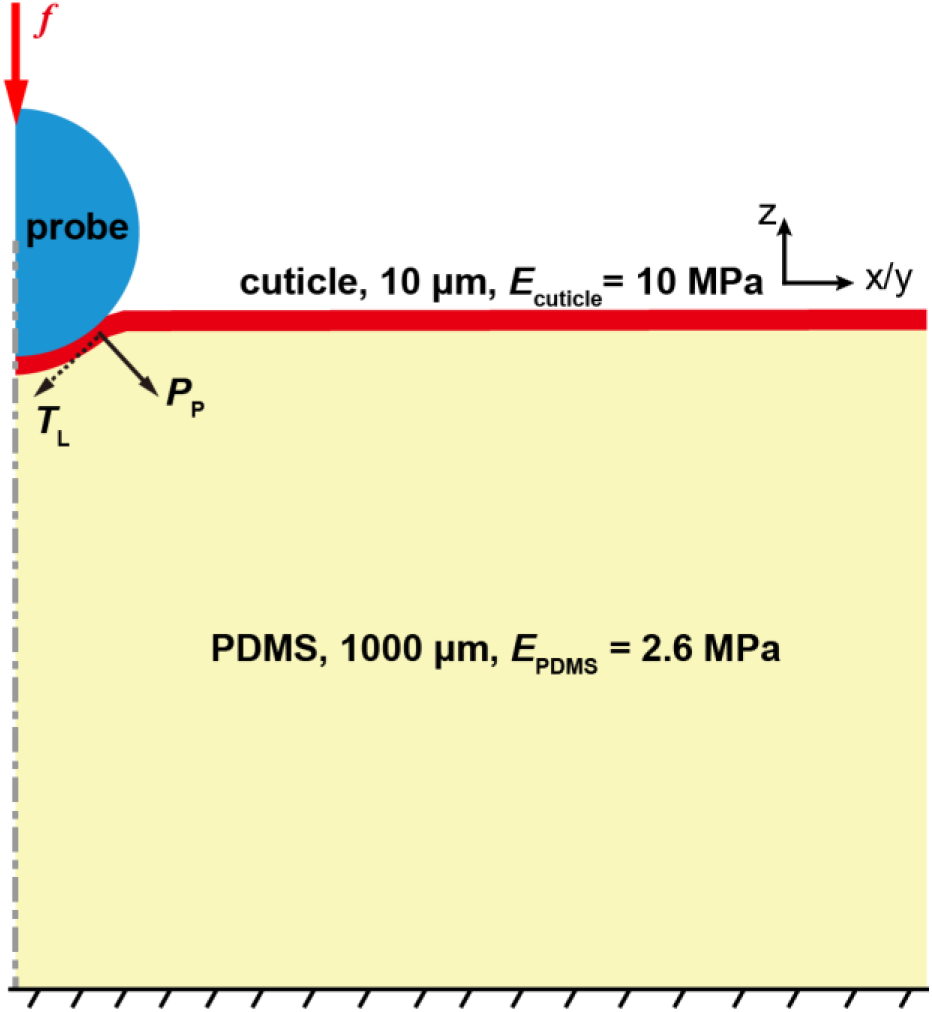
Cartoon schematics for the mechanical model used in the finite element analysis. *P*_P_, pressure perpendicular to the cuticle surface. *T*_L_, tension parallel with the cuticle surface.

**Movie s1**

The representative response of a c4da cell to a 4 mN force stimulus. The asterisk indicated the force application point and the white arrowhead indicated the soma.

